# High-throughput cryo-EM enabled by user-free preprocessing routines

**DOI:** 10.1101/2019.12.20.885541

**Authors:** Yilai Li, Jennifer N. Cash, John. J.G. Tesmer, Michael A. Cianfrocco

## Abstract

The growth of single-particle cryo-EM into a mainstream structural biology tool has allowed for many important biological discoveries. Continued developments in data collection strategies alongside new sample preparation devices heralds a future where users will collect multiple datasets per microscope session. To make cryo-EM data processing more automatic and user-friendly, we have developed an automatic pipeline for cryo-EM data preprocessing and assessment using a combination of deep learning and image analysis tools. We have verified the performance of this pipeline on a number of datasets and extended its scope to include sample screening by the user-free assessment of the qualities of a series of datasets under different conditions. We propose that our workflow provides a decision-free solution for cryo-EM, making data preprocessing more generalized and robust in the high-throughput era as well as more convenient for users from a range of backgrounds.

## INTRODUCTION

Single-particle cryo-electron microscopy (cryo-EM) is becoming a mainstream technique for structural biology (Kühlbrandt 2014). In the past few years, cryo-EM has seen a 20-40% year-to-year growth in structures deposited in the Protein Data Bank. This growth is due to continued developments in sample preparation (Cheng et al. 2018; Zivanov et al. 2018; Jain et al. 2012; Arnold et al. 2017; Darrow et al. 2019; Ravelli et al. 2019), data collection (Fernandez-Leiro and Scheres 2016; Lyumkis 2019), and algorithms for data processing (Scheres 2012; Zivanov et al. 2018; Punjani et al. 2017; Tegunov and Cramer 2019). These developments have greatly accelerated the speed of data collection for cryo-EM, and have also led to widespread adoption of users across a range of expertise, where experts represent a continually shrinking fraction of cryo-EM users.

With the fast pace of cryo-EM development, several challenges have emerged. First, with new imaging and sample preparation technologies, including the increased frame rate detectors, beam-image shift data collection (Cheng et al. 2018; Zivanov et al. 2018), and robotic sample preparation (Jain et al. 2012; Arnold et al. 2017; Darrow et al. 2019; Ravelli et al. 2019), a single cryo-EM instrument can easily generate 5,000-8,000 movies data per day. These technologies have enabled cryo-EM to become a more high-throughput technique. Second, although a number of improvements have been made in software development, cryo-EM data processing remains computationally expensive. High-performance computing (HPC) resources and GPUs are typically used (Michael A. Cianfrocco and Leschziner 2015; Baldwin et al. 2018). However, since each project requires multiple rounds of human trial and error in the preprocessing steps, these human-driven choices can slow down a project due to a lack of computing resources.

Third, cryo-EM frustrates many users because of its complexity in data processing. The manual and subjective decisions involved in solving a structure, such as the programs, parameters, and determination of good micrographs and good 2D class averages, can affect the final result significantly (Lawson and Chiu 2018). While an expert can make the correct decisions after a few trials, new users typically find it problematic to perform such monitoring and evaluations. Moreover, due to the variety of samples in the cryo-EM field, it is nearly impossible to create a general guideline for the new users to follow.

Despite the increasing throughput of cryo-EM data collection, the cumbersome nature of cryo-EM preprocessing slows scientists’ ability to ask biological questions from their dataset. For example, during cryo-EM sample screening, scientists may want to assess sample integrity or complex formation. However, in order to compare and contrast multiple grids, the scientist will have to manually interact with the data to perform movie alignment, particle picking, CTF estimation, and 2D classification. Modern cryo-EM needs a tool to streamline data quality assessment and data preprocessing automatically and robustly.

Many approaches have been proposed and developed to address these challenges. For example, Appion (Lander et al. 2009), cryoSPARC (Punjani et al. 2017), SPHIRE (Moriya et al. 2017), and RELION-3.0 (Zivanov et al. 2018; Fernandez-Leiro and Scheres, n.d.) provide preprocessing tools that can be stitched together into pipelines. Despite this ability, easy computation access to these remains an issue. To address the computation resource problem, COSMIC^2^ (M. A. Cianfrocco et al. 2017), a science gateway for cryo-EM, has been developed with the philosophy of bringing popular cryo-EM tools and resources to all scientists in the field, removing the practical limitations that accessing those resources would otherwise entail.

Many algorithms have also been developed to accelerate cryo-EM data preprocessing and minimize subjective decisions and tedious human annotations. Notably, deep learning, especially convolutional neural network (CNN), has greatly changed and improved the step of particle picking (Wagner et al. 2019; Bepler et al. 2019; Tegunov and Cramer 2019; Wang et al. 2016; Zhu, Ouyang, and Mao 2017; Zhang et al. 2019; Xiao and Yang 2017; Nguyen et al. 2019; Al-Azzawi et al. 2019). Nevertheless, the field still lacks a robust tool that will make decisions by evaluating the output from data preprocessing steps, so that human intervention can be removed, making an automatically streamlining workflow possible.

Here, we introduce several deep learning and image analysis tools for automated preprocessing and assessment of cryo-EM datasets. By connecting these tools with state-of-the-art data preprocessing algorithms, we make a general workflow that can achieve expert-level performance on a number of different cryo-EM datasets. Our workflow takes movies or motion-corrected micrographs as the input and outputs a particle stack that contains high-resolution particles that will be used in the following 3D reconstruction steps without any user decisions. Specifically, our workflow can automatically detect bad micrographs using *MicAssess*, determine the best parameters for particle picking and 2D classification, and identify the good class averages that can be used in 3D reconstruction using *2DAssess*. In the workflow, the subjective user decisions are replaced with statistical models based on the features extracted with image processing methods and convolutional neural networks, along with the expert knowledge. We believe that our automatic pipeline helps to establish a framework to accelerate data preprocessing and to perform data assessment at multiple levels in the high-throughput era of cryo-EM.

## RESULTS

### Overview of the Method

The current routine of cryo-EM data preprocessing consists of a number of subjective user decisions (**Fig. 1**). First, many users will manually go through all the motion-corrected micrographs to pick out the bad micrographs, and then select an estimated resolution threshold to remove the remaining bad micrographs based on the results of CTF estimation. Next, most particle pickers will require the users to manually pick a few particles, set the estimated particle diameter and determine the picking threshold before automatic particle picking. Then the particles will be extracted with the user-defined box size and pixel size used for 2D classification, where the users need to determine the class number and the diameter of the mask. Finally, the users need to select the good 2D class averages based on their own judgment, and the particles in the selected 2D class averages will be re-extracted and used in the downstream 3D reconstruction steps.

**Figure 1.**
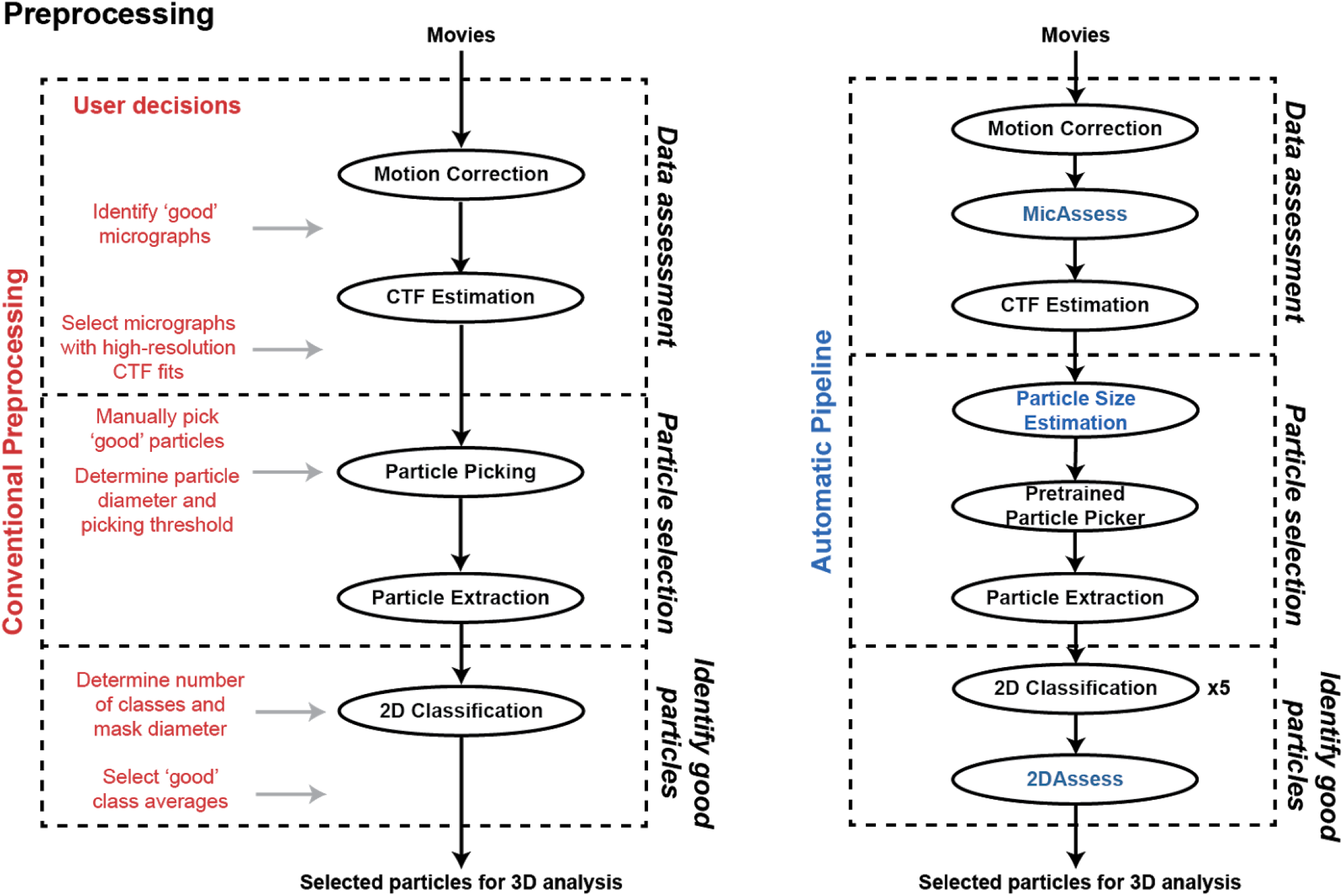
Conventional cryo-EM preprocessing vs. automatic preprocessing pipeline. Left panel: current workflow describing the preprocessing of cryo-EM datasets, with all the user decisions needed in red. Right panel: the automatic pipeline introduced in this paper. All user decisions are replaced by the new tools developed in blue.

Our general workflow streamlines the preprocessing steps to take either movies or motion-corrected micrographs as the input and output a stack of clean particles that can be used as the input in the subsequent 3D analysis (**Fig. 1**). During this process, we built statistical models in order to capture human decision-making during the preprocessing steps. Instead of developing new preprocessing tools and algorithms, our workflow takes advantage of these developments and provides evaluations so that expert-level decisions can be made automatically. We provide an overview of the method below.

#### *MicAssess:* Automatic micrograph assessment

First, we developed a tool that can assess the quality of motion-corrected micrographs even before CTF estimation: *MicAssess*. Unlike EMPIAR datasets, which consist of mostly usable micrographs, many real-world data generated from the microscopes are dirty and noisy. Researchers often undertake significant effort to manually eliminate bad micrographs to obtain a clean dataset to work within the downstream preprocessing steps. Although the difference between good and bad micrographs is unambiguous, it is still difficult to find a universal and robust criterion. Many scientists have been using the resolution outputs from CTF estimation for micrograph cleaning, however, there lacks a publicly accepted resolution cutoff, and there are still a number of bad micrographs that will make through using this metric for decision making.

Convolutional neural networks (CNN) are changing the field of computer vision as well as biology in recent years and have been widely applied to image classification, object detection, image segmentation, etc (Moen et al. 2019). In cryo-EM, a number of CNN-based particle picking models have been developed and widely used, including Warp (Tegunov and Cramer 2019), crYOLO (Wagner et al. 2019), and Topaz (Bepler et al. 2019). With the similar idea, we developed a CNN-based micrograph assessor, *MicAssess*. The architecture of *MicAssess* is described in **Fig. 2A**. Similar to many CNN models, our model consists of a feature extraction convolutional network and a classification network. For the feature extraction network, we used a standard ResNet34 (He et al. 2016), which is a deep and light-weighted fully convolutional residual network with 34 layers. Following the feature extraction, the convolutional network is the classification network, which consists of one fully connected (FC) layers with 512 nodes. Dropout layers with a 0.5 dropout rate and batch normalization are also applied, and leaky rectified linear unit (LReLU) is used as the activation function. Finally, the last layer uses a sigmoid function as the activation function and performs prediction, which is the probability that the input micrograph is considered as “good”.

**Figure 2.**
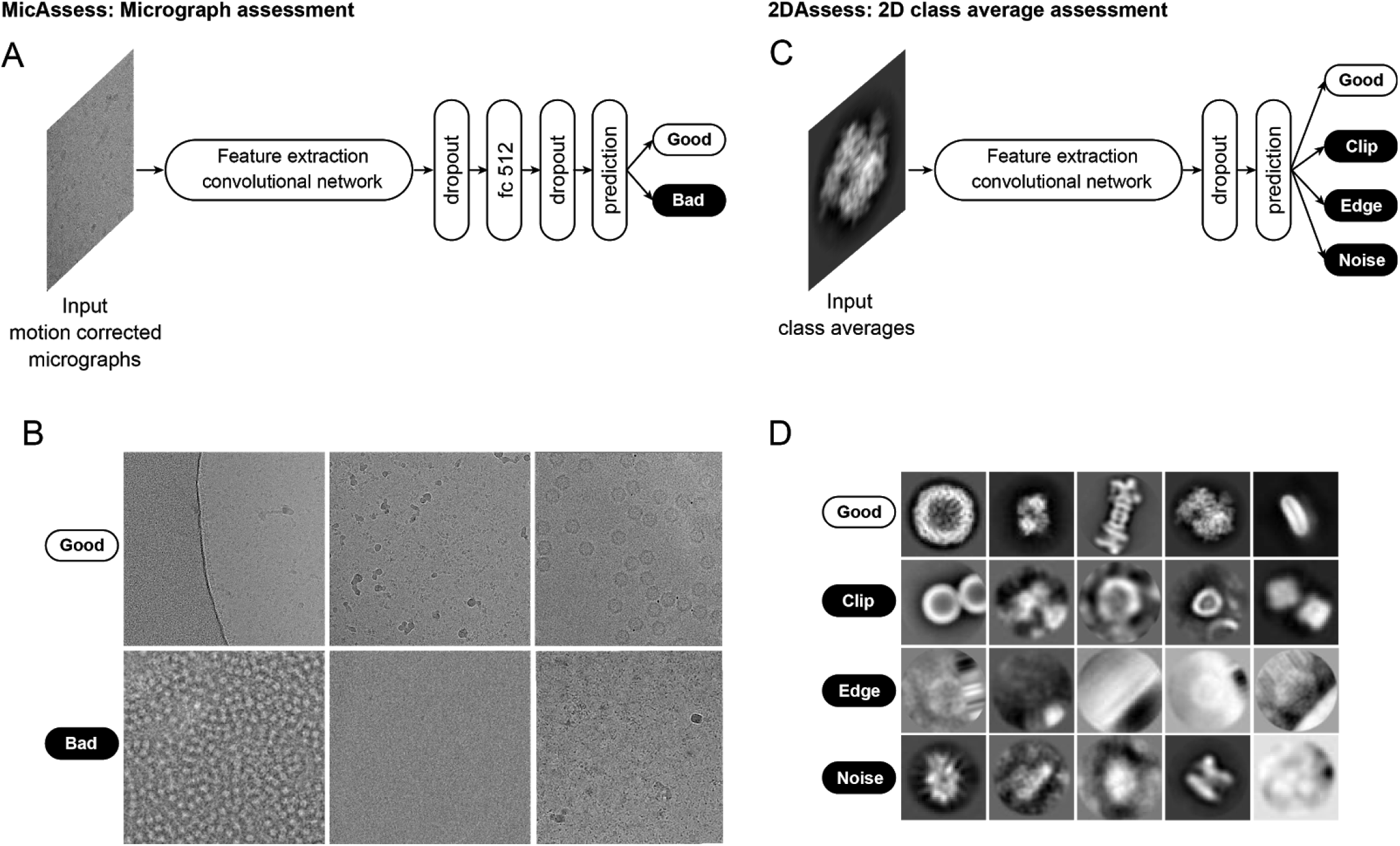
Deep learning-based tools for cryo-EM micrograph and 2D class average assessment. (A) The architecture of *MicAssess*. The motion corrected micrograph will be inputted to a feature extraction convolutional network (a standard ResNet34 in the paper), and after one dropout layer, one fully connected layer and another dropout layer, output the prediction of the micrograph. (B) Examples of the labeled good and bad micrographs in the training set. The good class contains partially good images, images with small or very large proteins, etc.. The bad class contains all different kinds of unusable micrographs, including micrographs that are empty or too dense, contaminated, or with protein aggregates. (C) The architecture of CNN-based model in *2DAssess*. The input class average image will be inputted to a feature extraction convolutional network (a standard ResNet50 in the paper), and after one dropout layer, output the prediction of the 2D class average to be one of the four classes. (D) Examples of the labeled 2D class averages in the good, clip, edge, and noise classes in the training set.

Most image classification problems are considered as supervised learning, which means that they need to be trained on labeled datasets. We have collected and manually labeled a total of 4,644 micrographs (2,372 good micrographs and 2,272 bad micrographs) from several EMPIAR datasets in addition to in-house datasets (**Table 1**). Our good micrograph dataset consists of proteins and complexes ranging from 50 kDa to 4 MDa (**Fig. 2B, upper row**), while our bad micrograph dataset consists of a variety of unusable micrographs including micrographs that are either empty or too dense, contaminated, or with protein aggregates (**Fig. 2B, lower row**). The dataset was randomly split into a training set (80 %) and a validation set (20 %). Data augmentation was applied before training to increase the amount of training data and reduce overfitting. The trained model was evaluated on the validation set, and an accuracy of about 97% was achieved. A detailed description can be found in the Methods section.

**Table 1.** Sources of the micrographs in the training and validation dataset used for *MicAssess*.

| Particle Name | EMPIAR ID |
| --- | --- |
| 26S Proteasome | EMPIAR-10072 |
| AAV | EMPIAR-10202 |
| Ribosome | EMPIAR-10077 |
| Rag complex | EMPIAR-10049 |
| NOMPC | EMPIAR-10093 |
| GluDH | EMPIAR-10217 |
| RNA PolIII | EMPIAR-10190 |
| Spliceosome | EMPIAR-10160 |
| In-house dataset - 160 kDa | N/A |
| In-house dataset - 480 kDa | N/A |
| In-house dataset - 180 kDa | N/A |
| In-house dataset - 168 kDa | N/A |
| In-house dataset - 80 kDa | N/A |

To test the effectiveness of *MicAssess*, we analyzed a published dataset collected by our lab on the Phosphatidylinositol 3,4,5-trisphosphate (PIP_3_)-dependent Rac exchanger 1 (P-Rex1) (Cash et al. 2019). This dataset contains 6,736 micrographs and is a combination of untilted and tilted series. Importantly, the training data in *MicAssess* did not include any P-Rex1 micrographs. As a comparison, we also classified the micrographs using the CTF maximum resolution outputs from CTFFIND4, with determination thresholds being 4 Å for untilted micrographs and 10 Å for tilted micrographs. To quantify the performance of both CTF-based micrograph cleaning and *MicAssess*, we manually labeled the total 6,736 micrographs and used the labels as the “ground truth” with which to compare.

A comparison of CTF maximum resolution cutoff to the ground truth highlighted a number of discrepancies. As is typical, the distribution of CTF maximum resolution values for tilted or untilted micrographs does not show a bimodal distribution. (**Fig. 3A**). Therefore, even though 4 Å and 10 Å resolution cutoff thresholds are considered reasonable, such numbers are not obvious from the distribution of the data, but rather arbitrary. Compared to human-labeled “ground truth”, CTF-based micrograph cleaning reached an overall accuracy of 77.5% (**Fig. 3B**). This indicates that while CTF maximum resolution is a convenient method to remove bad micrographs, there is room for improvement in order to obtain more accurate micrograph assessment.

**Figure 3.**
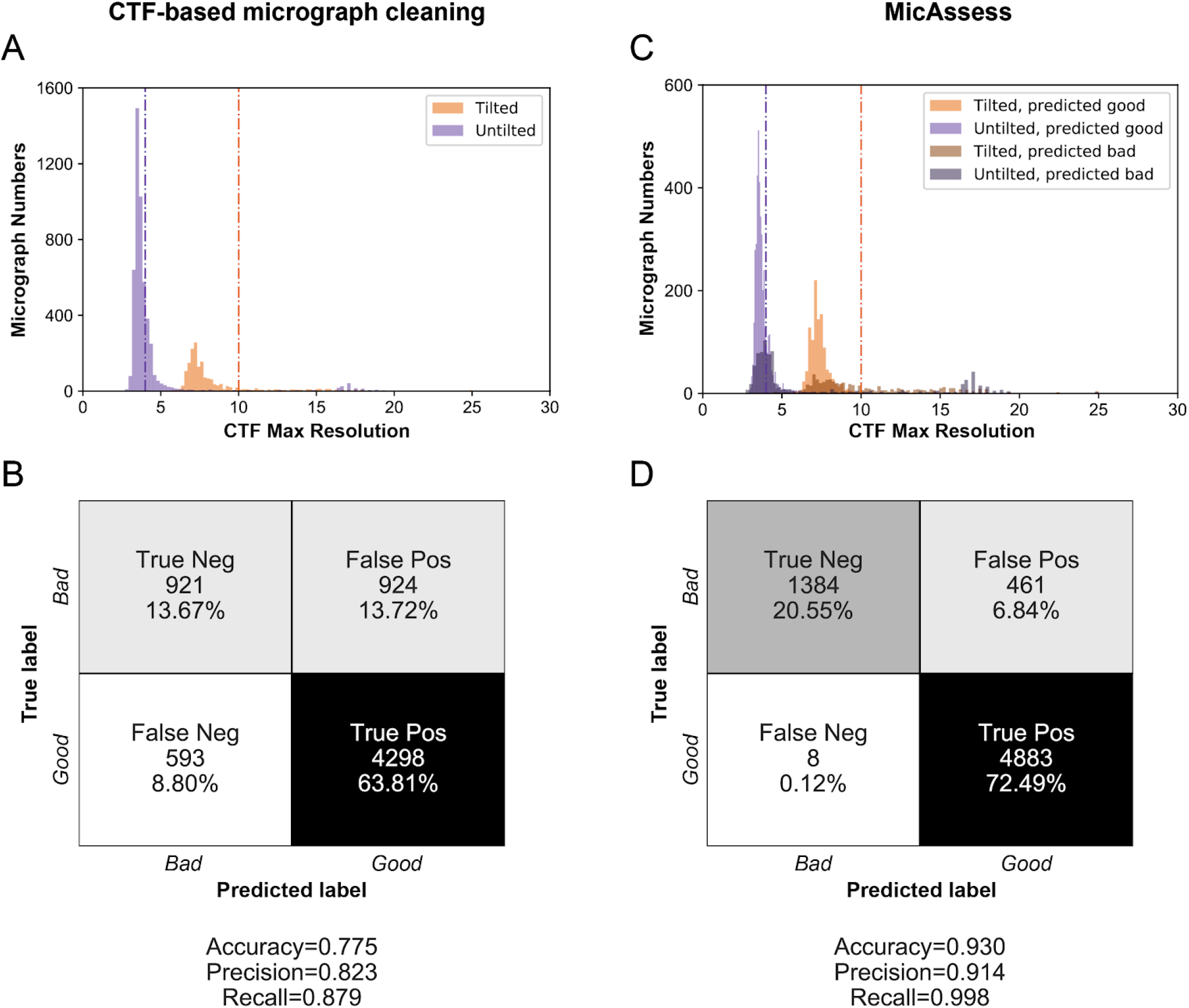
*MicAssess* performs equivalently to CTF resolution cutoff on micrograph assessment. (A) Histograms of the CTF maximum resolutions outputted by CTFFIND4 of the test set. Vertical lines indicate the selected hard thresholds for tilted and untilted micrographs (4 Å and 10 Å respectively). Micrographs higher than the thresholds are considered as bad. (B) Confusion matrix and evaluation metrics for CTF resolution threshold vs. human assessment on P-Rex1:Gβγ dataset. (C) Histograms of the CTF maximum resolutions outputted by CTFFIND4 of the test set, color labeled according to the predictions by *MicAssess*. Vertical lines indicate 4 Å and 10 Å respectively. (D) and evaluation metrics of *MicAssess* on the P-Rex1 test set.

Compared to CTF maximum resolution, *MicAssess* showed higher accuracy for identifying both good and bad micrographs. To highlight the power of *MicAssess*, *MicAssess* was also able to correctly classify many bad micrographs with < 4 Å CTF maximum resolutions (**Fig. 3C**). Such micrographs will not be captured by the CTF-based micrograph cleaning approach. Overall, *MicAssess* found 1,388 bad micrographs (**Fig. S2**) and had an accuracy of 93.0%, with a notably very low false-negative rate (0.12%) (**Fig. 3D**). In other words, only 8 good micrographs were misclassified to the bad category.

This analysis indicates the *MicAssess* performs nearly as well as human assessment for the P-Rex1 test dataset. More importantly, *MicAssess* does not need any arbitrary threshold, and both tilted and untilted micrographs were predicted with the exact same procedure, providing a completely “hands-off” tool for micrograph assessment, which enables the automatic cryo-EM data preprocessing and assessment at the very beginning.

#### Automatic particle diameter estimation

Since our workflow aims for decision-free preprocessing, the suitable particle picker should not need any human picking beforehand. Therefore, any template-based particle picker or CNN-based particle picker that needs to be trained on manually prelabeled particles cannot be used in the workflow. Fortunately, we are able to use the general model of crYOLO (Wagner et al. 2019), which is a CNN-based particle picker pretrained on a number of EMPIAR and in-house datasets. The two parameters needed for particle picking in crYOLO are box size and threshold.

Optimally, the box size should be the size of the particle. Since this information is usually unclear for a new protein, our workflow will first perform particle picking on a subset of the micrographs with different box sizes. The picked particles will be extracted, low pass filtered and averaged without any alignment. We then find the edge of this averaged image using a Canny edge detector, and the size of the particle is determined based on the edge detected and is dilated by an empirical factor of 1.5 (**Fig. S3**). After that, the workflow uses crYOLO to pick the particles from all micrographs. The threshold parameter controls the strictness of the decision of a particle. The workflow uses a very low threshold of 0.1 since many false positives can be removed in the following 2D classification step.

#### *2DAssess*: Automatic selection of good 2D class averages

After particles are picked and extracted from micrographs with CTF information, particles are subjected to 2D classification, whereby good 2D averages are identified using *2DAssess*. Similar to the micrograph classifier, our CNN-based classifier model (**Fig. 2C**) also requires a labeled dataset for training. We have obtained the 2D class averages from ten different datasets from a range of diameters use in 2D classification, providing 2D averages for optimal masks, masks that are too tight, and masks that are too large (**Table 2**).

**Table 2.**
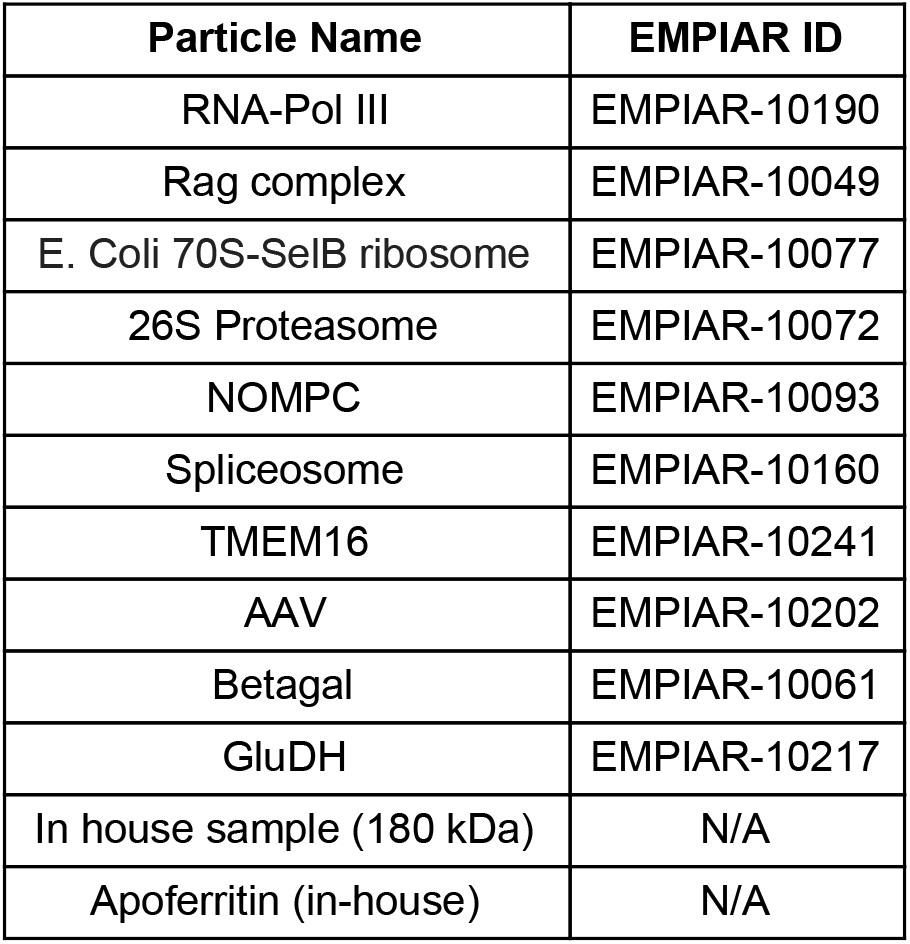
A full list of the 2D class averages in the training and validation dataset used for *2DAssess*.

The 2D class averages are preprocessed and labeled in four different classes (**Fig. 2D**): good, clip, edge, and noise. The good class includes all the good class averages that will be selected and used in the downstream processing steps. The clip class includes the class averages that are clipping the neighboring particles, usually a sign that the diameter is too large. The edge class includes the class averages with “barcode” like patterns, which means that some particles are on the edge of the micrograph or the carbon. The noise class includes all the other bad class averages that are not covered by the clip and edge classes, which contains pure noise, over-aligned, and low-resolution class averages. The dataset was downsampled to account for the class imbalance and then randomly split into a training set (80 %) and a validation set (20 %). We noticed that when the diameter of the mask becomes large, one class average might contain two particles. The CNN-based classifier failed to detect this and would misclassify such 2D class averages to the “good” class. To prevent this, we checked the saliency map (Hou and Zhang 2007) of the 2D class averages in the predicted “good” class, and re-classify the class averages with two or more objects to the correct “clip” class. The combination of the CNN-based classifier and the saliency map check made up the complete 2D class average assessor, which we named it as *2DAssess*.

To further enrich the number of good class averages, we used deep convolutional generative adversarial networks (DCGAN) (Radford, Metz, and Chintala 2015) to generate artificial good class averages using the true good class averages in the training set. We then carefully selected 66 artificial good class averages generated by DCGAN (**Fig. S4**) and added them to the training set. Although the selected images are not from 2D class averages of real proteins, they will most likely be labeled as good class averages without any prior knowledge of the protein. Adding these DCGAN generated images as a data augmentation approach improves the generalizability of the classifier when the good 2D class average samples are limited. Some simple data augmentation (elaborated in the methods section of the paper) was applied in training and validation. The precision and recall of each class for the validation set are reported in **Table 3**. Notably, the good class reached a precision of 94% and a recall of 97%.

**Table 3.**
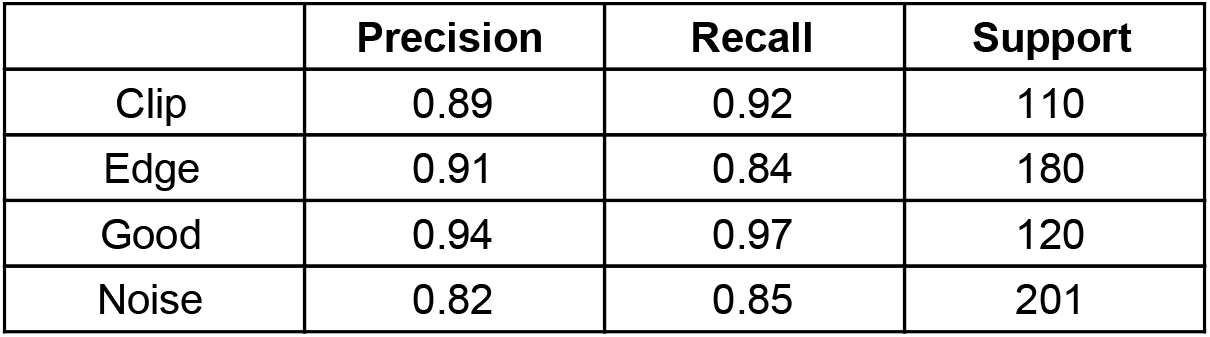
Precisions, recalls and the number of supports of each class in the validation set of *2DAssess*.

### Testing on EMPIAR Datasets

#### T20S proteasome (EMPIAR-10025)

First, we tested our workflow on a subset of the published T20S proteasome cryo-EM dataset (EMPIAR-10025) (Campbell et al. 2015) (**Fig. 4**). This subset contains 87 micrographs, all of which were all being classified as good by *MicAssess*. Subsequently, the diameter was estimated to be 195 Å. Using this diameter, crYOLO picked 52,153 particles that were used to search a range of diameters during 2D classification (**Fig. 4A & B**). For each diameter used in RELION 2D classification, *2DAssess* was used to estimate the number of good particles. Finally, comparison across all diameters used in 2D classification indicated that the best diameter for T20S was 195 Å (**Fig. 4B**). For the 195 Å diameter, the good 2D class averages selected by *2DAssess* had a 100% prediction accuracy (**Fig. 4C**), correctly identifying all good and bad 2D averages.

**Figure 4.**
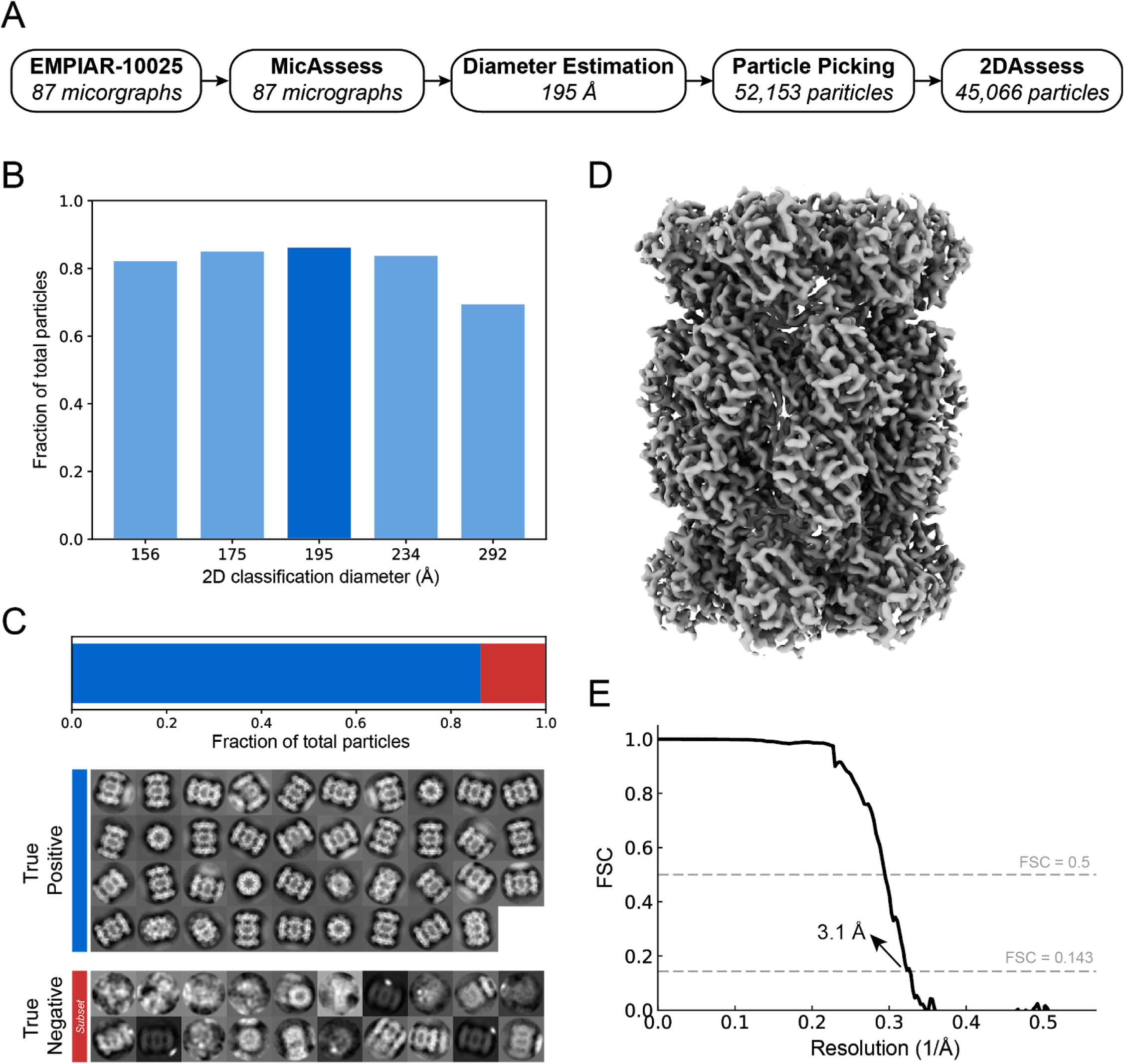
High-resolution cryo-EM structure of T20S proteasome from automatic preprocessing pipeline. (A) Overview of the intermediate results of automatic pipeline on EMPIAR-10025 dataset. (B) Histogram showing the fractions of the good particles identified by the pipeline with different diameter used in 2D classification. The diameter with the most good particles (195 Å) is selected (darker blue) to be the best diameter, and the corresponding 2D classification result is used to output the final particle stack. (C) *2DAssess* achieves 100% prediction accuracy on the EMPIAR-10025 dataset. All the good 2D averages (86.4% of the picked particles) and a subset of the bad 2D averages predicted by *2DAssess* are shown. (D) 3D electron density volume using the particle stack outputted by the pipeline as the input for 3D reconstruction steps. (E) FSC curve of the electron density map in panel C, showing a resolution of 3.1 Å.

Using the stack of particles associated with good averages, we then performed 3D refinement to obtain a 3.1 Å structure of the T20S proteasome (**Fig. 4D & E**). This structure demonstrates that the automatic preprocessing pipeline provided a high-resolution stack of particles of T20S without user intervention.

#### Hemagglutinin (HA) trimer (EMPIAR-10175)

After successfully analyzing T20S, we next wanted to try a more challenging sample. To this end, we selected the influenza hemagglutinin trimer (HA trimer) dataset (EMPIAR-10175) (Noble et al. 2018) due to its extreme orientation differences: end-on views have a diameter of 55Å whereas the side-on views have a diameter of 140Å. After running *MicAssess* on 1,099 micrographs, *MicAssess* identified 205 micrographs as bad (examples are shown in **Fig. S5**), and the rest of 894 micrographs were preprocessed by the downstream pipeline. After 2D classification, the best diameter to be used in 2D classification was selected to be 150 Å (**Fig. 5A & B**). The good and bad class averages were all correctly classified by *2DAssess* (**Fig. 5C**).

**Figure 5.**
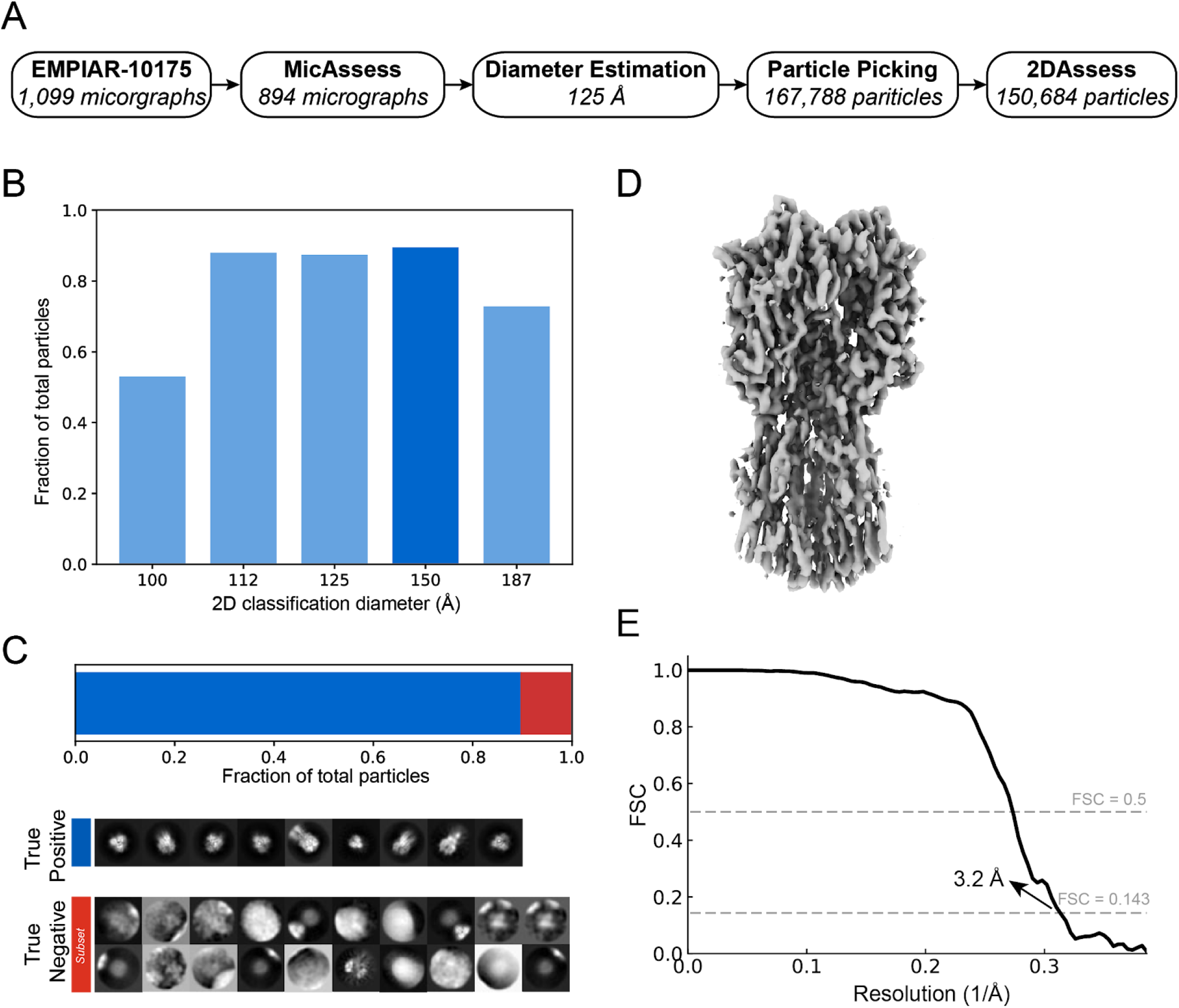
High-resolution cryo-EM structure of HA trimer from automatic preprocessing pipeline. (A) Overview of the intermediate results of automatic pipeline on EMPIAR-10175 dataset. (B) Histogram showing the fractions of the good particles identified by the pipeline with different diameter used in 2D classification. The diameter with the most good particles (150 Å) is selected (darker blue) to be the best diameter, and the corresponding 2D classification result is used to output the final particle stack. (C) *2DAssess* achieves 100% prediction accuracy on the EMPIAR-10175 dataset. All the good 2D averages (89.8% of the picked particles) and a subset of the bad 2D averages predicted by *2DAssess* are shown. (D) 3D electron density volume using the particle stack outputted by the pipeline as the input for 3D reconstruction steps. (E) FSC curve of the electron density map in panel C, showing a resolution of 3.2 Å.

Using the output stack of good particles, we performed a 3D refinement with the selected 150,684 particles. This allowed us to determine a structure at 3.2 Å resolution (**Fig. 5D & E**), comparable to what was published previously for HA trimer (Noble et al. 2018). This structure confirmed that the automatic pipeline is capable of handling datasets of varying size and shape, setting the stage for real-world data analysis.

### Analysis of real-world data

#### Aldolase

To extend our preprocessing pipeline, we analyzed an in-house collected aldolase dataset. This dataset contains 1,118 micrographs, in which 1,075 micrographs were predicted as good by *MicAssess*. The examples of bad micrographs being selected by *MicAssess* are shown in **Fig. S6**. After estimating the particle diameter, the 2D classification showed an optimal mask diameter of 108 Å (**Fig. 6A & B**). *2DAssess* correctly predicted all the good class averages. In this dataset, there were two falsely identified good averages that were actually bad, which only accounted for 1.53% of the total particles (**Fig. 6C**).

**Figure 6.**
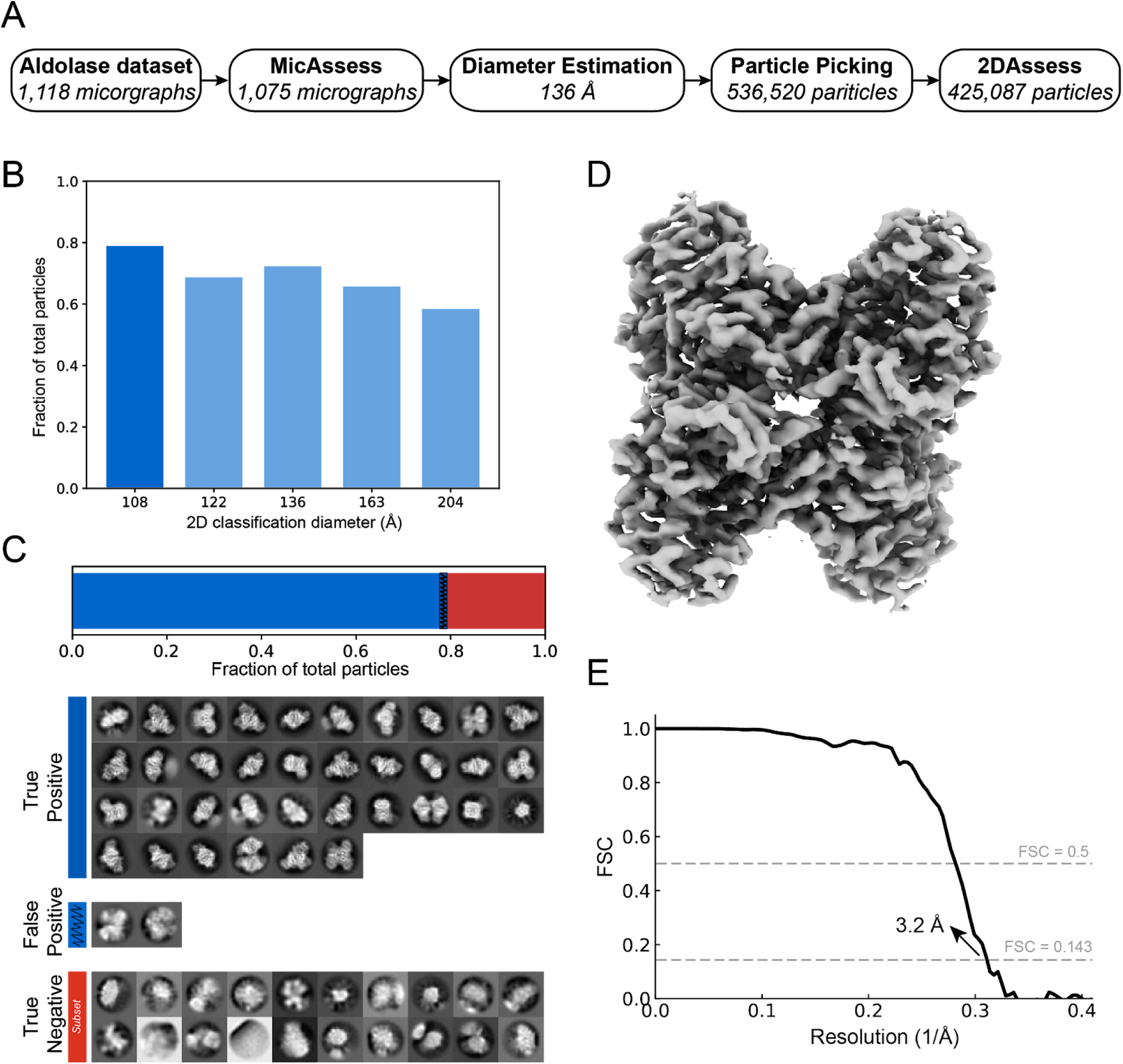
High-resolution cryo-EM structure of aldolase from automatic preprocessing pipeline. (A) Overview of the intermediate results of automatic pipeline on the aldolase dataset. (B) Histogram showing the fractions of the good particles identified by the pipeline with different diameter used in 2D classification. The diameter with the most good particles (108 Å) is selected (darker blue) to be the best diameter, and the corresponding 2D classification result is used to output the final particle stack. (C) *2DAssess* achieves very high prediction accuracy on the aldolase dataset. All the good 2D averages (79.2% of the picked particles) and a subset of the bad 2D averages predicted by *2DAssess* are shown. The two false positives (blue shaded) only account for 1.53% of the total picked particles. (D) 3D electron density volume using the particle stack outputted by the pipeline (including the false positives) as the input for 3D reconstruction steps. (E) FSC curve of the electron density map in panel C, showing a resolution of 3.2 Å.

Using the particle stack generated by the pipeline (including all of the false positives), we performed a 3D refinement to obtain a final structure of aldolase at 3.2 Å (**Fig. 6D & E**). This demonstrates that the preprocessing pipeline successfully handles more realistic data, as expert users also determine a structure to the same resolution.

#### P-Rex1 - a sample screening case study for high-throughput cryo-EM

Finally, in order to demonstrate the effectiveness of the pipeline, we automatically analyzed multiple datasets to simulate a sample screening experiment. The datasets we used were collected from six cryo-EM sessions of P-Rex1 under different conditions (**Fig. 7A**), including apo P-Rex1 on different types of grids (18sep06b and 18sep28b), with different additives (18jan09b and 18jan09d), and with a binding partner Gβγ at different concentrations (18jul14a and 18jan18c). The goal of this sample screening case study is to verify that our pipeline provides a robust and user-free approach for automatic data quality assessment at different levels considering that only one dataset (18jan18c) is amenable for high-resolution cryo-EM (Cash et al. 2019).

**Figure 7.**
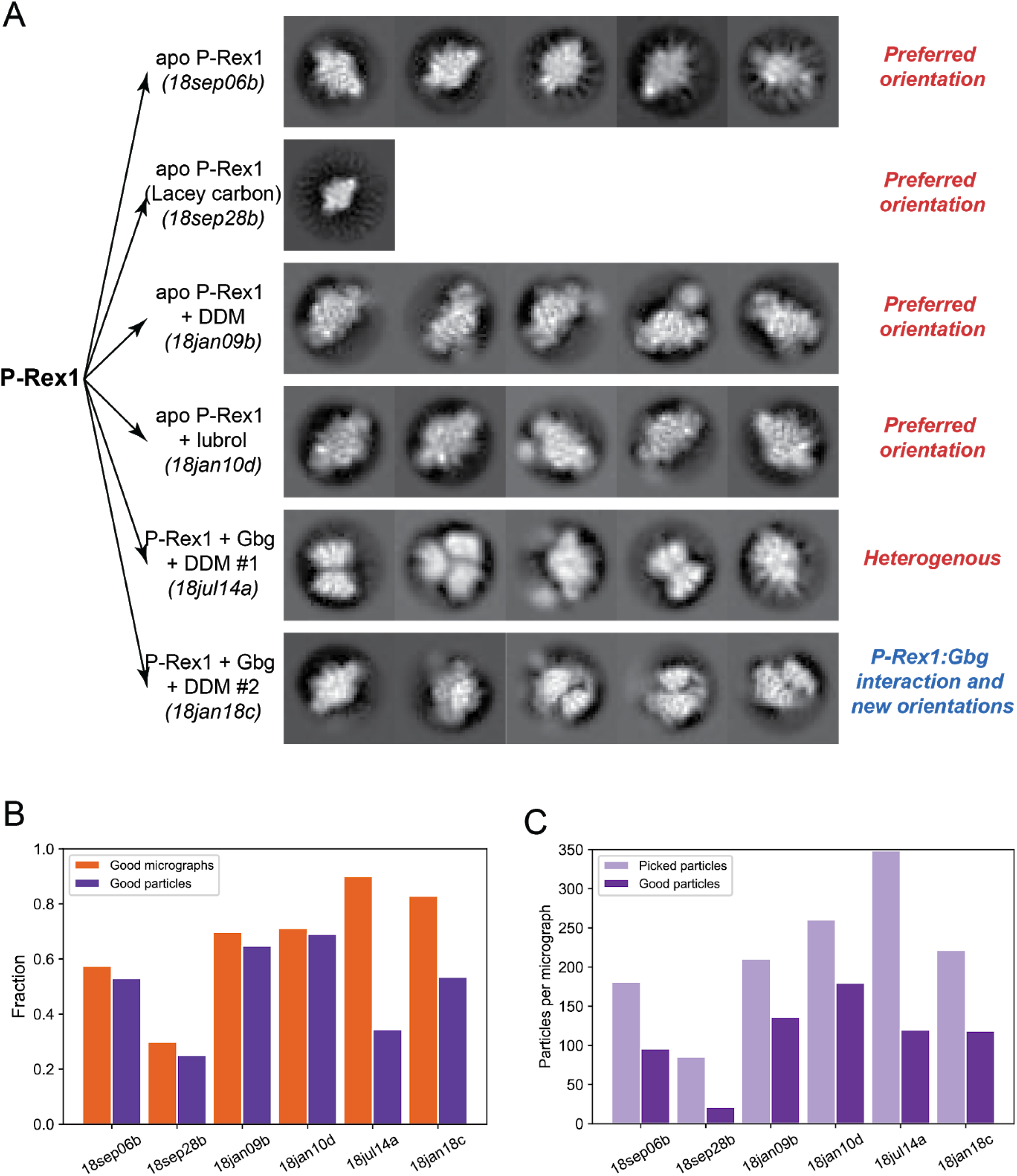
Automatic analysis of multiple P-Rex1 cryo-EM data sets to assess sample quality. (A) The six datasets analyzed by the automatic pipeline in this case study, including different sample preparations, different additives and whether a binding partner was added. 2D class averages were predicted by 2DAssess and the five good and representative 2D class averages for each dataset are shown for assessment. (B) Fractions of the good micrographs in all the micrographs (orange) and fractions of the good particles outputted by the automatic pipeline in all the picked particles (purple) for each dataset. (C) The numbers of picked particles (brown) and the numbers of good particles outputted by the automatic pipeline (blue) for each dataset.

All six datasets were analyzed with the pre-defined automatic pipeline, where no user input was required other than microscope settings. The outputs of the automatic pipeline were the 2D class averages selected by *2DAssess* for each dataset (**Fig. 7A & Fig. S7**). The datasets were assessed at different levels, from the micrographs to the 2D class averages, throughout the pipeline. At the first step, *MicAssess* quickly captured that one of the datasets, 18sep28b, contained mostly bad micrographs (70%) (**Fig. 7B**). All the other five datasets contained mostly good (above 50%) micrographs (**Fig. 7B**). The particle picker picked 170-350 particles per micrograph for all five datasets, except 18sep28b, which only had an average of 85 picked particles per micrograph, confirming the bad quality of this dataset (**Fig. 7C**). After 2D classification, the class averages were classified by *2DAssess*, where we found that four datasets have over 50% of the picked particles to be good particles outputted by the automatic pipeline (**Fig. 7B**), and there were 100~200 good particles per micrograph (**Fig. 7C**).

Although many of the datasets showed a promising statistics of good micrograph and good particle fractions, the good 2D class averages selected by *2DAssess* revealed that apo P-Rex1 alone and with additives had a very strong preferred orientation on the cryo-EM grids (**Fig. 7A**). On the other hand, one of the datasets of P-Rex1 with Gβγ (18jul14a) exhibited a sample heterogeneity, where we found Gβγ oligomers in the good 2D class averages (**Fig. 7A**), indicating that the concentration of Gβγ added was too high. Finally, the 2D class averages output by the automatic pipeline from the last dataset (18jan18c) showed the P-Rex1 and Gβγ interactions, and new orientations were also seen as a result (**Fig. 7A**). This case study demonstrated that our automatic preprocessing pipeline is an objective, fully automatic approach for sample screening for high-throughput cryo-EM.

## DISCUSSION

Cryo-EM is on the verge of becoming a high-throughput technique. This new era requires consistent and reproducible methods to assess and preprocess the micrographs directly from the microscopes in a timely manner. Our workflow provides a robust way to assess and preprocess cryo-EM data automatically without any user intervention, and it takes advantage of pre-existing software and preprocessing algorithms. We maintained the flexibility to incorporate any preprocessing algorithms, as long as no subjective user decisions are required. Therefore, instead of competing with the state-of-the-art software packages, our workflow uses the deep-learning-based assessment tools we developed and provides a platform to streamline all the preprocessing steps. To our knowledge, this is the first fully automatic and generic workflow for cryo-EM data preprocessing.

As the initial step in our workflow, it is important that *MicAssess* can efficiently identify most bad micrographs, but keeping ideally all the good micrographs. Therefore, *MicAssess* was tuned to tolerate more false positives, reducing the risk of a good micrograph being misclassified. The P-Rex1 benchmark result showed that it can effectively identify most of the bad micrographs from a big real-world dataset. Furthermore, *MicAssess* also has the potential to be incorporated into the data acquisition step. With the new K3 camera, which can collect as fast as 8,000 movies per day, it is impossible to manually assess the quality of the newly collected micrographs. *MicAssess* provides a way to assess these micrographs on the fly even before CTF estimation so that the user can get real-time feedback on the qualities of the micrographs.

In our workflow, we only used *2DAssess* to predict whether a class average is good or bad, but it can predict four different classes (clip, edge, good, and noise), which contains a lot more information. For example, a large percentage of particles being classified as “clip” usually indicates the mask diameter is too large because neighboring particles are being included in some 2D class averages. This gives the *2DAssess* the potential to improve 2D classification by performing automatic diameter searching. Specifically, since most 2D classification algorithms are iterative, intermediate 2D class averages are generated after each iteration. It is possible to apply *2DAssess* on the 2D class averages in the early iterations and use the outputted predictions to guide the automatic diameter searching.

Given that *MicAssess* and *2DAssess* are deep-learning based models, both models will continue to improve with more representative training data. Moreover, as deep learning models, these tools can be tuned for specific samples, users, or facilities to aid in sample assessment. Sample tuning could be extended into other parts of the pipeline, including particle picking and, likely, 3D analysis. Further work in this area stands to help streamline initial phases of cryo-EM data processing.

An important aspect of our pipeline centers on creating a workflow that does not depend on user-defined thresholds. These thresholds are typically CTF maximum resolution and particle picking thresholds, but could also apply to how 2D class averages are selected. By developing statistical tools to assess the data, we developed tools that more closely mirror user-based decisions, instead of fixed-value thresholds.

While this pipeline provides an important first step for automated pre-processing, there remains room for improvement. Namely, we continued to use 2D classification as a tool in order to measure particle quality, where belonging to ‘good’ class averages was a criteria for subsequent 3D analysis. Future work into particle sorting stands to provide a quick readout of particle quality to enable faster preprocessing routines.

Overall, this work demonstrates that user-free preprocessing is capable to perform in a manner comparable to that of an expert. Future work may extend into automated 3D analysis to enable cryo-EM users to quickly analyze multiple datasets in parallel.

## Acknowledgments

We would like to thank members of the Cianfrocco Laboratory for discussions related to this work. We would also like to thank Dr. Min Su for his help in discussing tilted CTF analysis. This work is supported by NSF-DBI-ABI 1759826 (Y.L. & M.A.C.) and R01-CA-221289 (J.N.C., J.J.G.T, & M.A.C.). The research reported in this publication was supported by the NIH under award number S10OD020011.

## Data accessibility

Cryo-EM structures have been deposited to the EMDB under accession codes EMDXXXX (T20S), EMDXXXX (HA Trimer), and EMDXXXX (Aldolase). All datasets are deposited to EMPIAR under XXXXX…

## Software availability

Software tools capable of running *MicAssess* and *2DAssess* will be available at https://github.com/cianfrocco-lab/Automatic-cryoEM-preprocessing. The preprocessing pipeline will also be incorporated into the freely available COSMIC^2^ science gateway: https://cosmic2.sdsc.edu:8443/gateway/.

## Competing financial interests

The authors do not have any competing financial interests for work in this manuscript.

## METHODS

### Automatic pipeline

#### MicAssess

Motion corrected micrographs were low-pass filtered and cropped to downscale to the network input image size of 494×494. Micrographs were then normalized to a mean of zero. A circular mask with diameter 494 pixels was applied to each micrograph, and then rotations and flipping were applied randomly in the training and validation dataset. The model was a 34-layer ResNet connected to two fully connected layers with leaky ReLU as the activation function and 0.5 dropout rate. The final predicting layer used a sigmoid function as the activation function. The loss function used was the binary cross-entropy loss. We used the ADAM optimizer with 0.0001 learning rate in training for optimization. In the real prediction, in order to tolerate more false positives than false negatives, we set the threshold as 0.1 (i.e. only micrographs with probabilities of being good lower than 0.1 will be classified as bad).

#### CTF estimation

CTF estimation is performed using CTFFIND4 (Rohou and Grigorieff 2015), with all the parameters, including pixel size, spherical aberration, magnification, and voltage, are related to the experiment given earlier.

#### 2D classification

Picked particles were scaled to about 3 Ångstrom/pixel and extracted using RELION3 (Zivanov et al. 2018). After that, all the particles will be processing with 2D classification in RELION3. The workflow uses the maximum class number, 200, for the best performance in the sacrifice of speed. Multiple 2D classification jobs for one dataset will be submitted, with different diameters of the mask, ranging from 0.5 to 2 times the particle size estimated earlier.

#### 2DAssess

Training and validation data consist of the RELION (Zivanov et al. 2018) outputs of 2D classification from 12 different datasets (**Table 3**). The EMPIAR datasets were preprocessed by the pipeline, and the outputted 2D class averages were manually labeled to the correct classes. Classes with significantly more samples were downsampled to eliminate the possible problems caused by class imbalance. The final dataset has 527, 550, 898, 1002 images for good, clip, edge, and noise classes respectively, and was randomly split into a training set (80 %) and a validation set (20 %).

Given that the output averaged images from RELION (Zivanov et al. 2018) already contained a mask with diameter *d*, we cropped all average images to remove mask edges. To do this, we first cropped the images to size *d*x*d* which only keep the centers of the images. Images were then normalized to a mean of zero, and resized to 256×256 using Lanczos resampling. Random rotations and flipping were applied in the training and validation dataset.

We used a simple DCGAN (Radford, Metz, and Chintala 2015) model to artificially generate images that belong to the good class as a data augmentation approach. The training data used for DCGAN is the 527 images in the good class. The generator of DCGAN was a convolutional neural network implementing upsampling convolutions, organized as input (100-d) -> transpose conv3×3 1024-d, stride 2, batch normalization, ReLU activation -> transpose conv1×1 1024-d, stride 1, batch normalization, ReLU activation -> transpose conv3×3 512-d, stride 2, batch normalization, ReLU activation -> transpose conv1×1 512-d, stride 1, batch normalization, ReLU activation -> transpose conv3×3 256-d, stride 2, batch normalization, ReLU activation -> transpose conv3×3 256-d, stride 2, batch normalization, ReLU activation -> transpose conv3×3 1-d, stride 1, tanh activation -> generated image. The discriminator of DCGAN was a simple convolutional neural network, organized as input -> conv3×3 32-d, stride 2, batch normalization, leaky ReLU activation, dropout rate 0.25 -> conv3×3 64-d, stride 2, batch normalization, leaky ReLU activation, dropout rate 0.25 -> conv3×3 128-d, stride 2, batch normalization, leaky ReLU activation, dropout rate 0.5 -> conv3×3 128-d, stride 2, batch normalization, leaky ReLU activation, dropout rate 0.5 -> fully connected layer with a single output with sigmoid activation. 10,000 epochs were used in training and only the images generated from the last 2,000 were saved. We then carefully selected 66 images and added them to the training set. All the selected images generated by DCGAN are shown in **Fig. S4**.

The CNN-based classifier failed to correctly classify class averages containing two particles, which is a situation that occurs when the 2D classification mask is too large. Therefore, we confirmed that all images predicted to be in the good class did not have two particles by calculating a saliency map of the 2D class averages. A saliency map is a representation of an image that can highlight the unique features of the image. In our application, we calculated the saliency map with the spectral residual approach and based on the object detected by the saliency map, we checked 1) the number of the object, and 2) whether the center of mass of the detected object is around the center of the image. Only the 2D class averages with one centered object detected will pass this saliency map check. The other class averages, with either more than one object or the object, are typically not well centered (usually due to the case that there are more than one particle but the particles are too close to be differentiated by the saliency map), will be moved to the correct clip class.

The number of the good particles that belong to the good 2D class average groups are calculated across all the diameters used in the 2D classification jobs, and the diameter with the best particles is being selected as the best diameter.

### Single-particle analysis

#### T20S

##### 3D refinement

After the preprocessing pipeline, 45,066 particles were re-extracted to a pixel size of 0.88 Å/pixel with a box size of 390 Å. Using EMD-6287 as an initial model, we performed a 3D refinement in RELION-v3.0 (Zivanov et al. 2018) using D7 symmetry to obtain a structure at 3.1 Å resolution and B-factor of −103 Å^2^.

#### HA Trimer

##### 3D refinement

After the preprocessing pipeline, 150,684 particles were re-extracted to a pixel size of 1.275Å/pixel with a box size of 250Å. Using EMD-7792 as an initial model, we performed homogenous 3D refinement in cryoSPARC v2.11.2-live_privatebeta using C3 symmetry to obtain a structure at 3.2 Å resolution and a B-factor of −151 Å^2^.

#### Aldolase

##### Sample preparation

Pure aldolase isolated from rabbit muscle was purchased as a lyophilized powder (Sigma Aldrich) and solubilized in 20 mM HEPES (pH 7.5), 50 mM NaCl at 1.6 mg/ml. Sample as dispensed on freshly plasma cleaned UltrAuFoil R1.2/1.3 300-mesh grids (Electron Microscopy Services) and applied to grid in the chamber of a Vitrobot (Thermo Fisher) at ~95% relative humidity, 4°C. The Sample was blotted for 4 seconds with Whatman No. #1 filter paper immediately prior to plunge freezing in liquid ethane cooled by liquid nitrogen.

##### Cryo-EM data acquisition

Data were acquired using the Leginon automated data-acquisition program (Suloway et al. 2005). Image preprocessing (frame alignment and CTF estimation) were done using the Appion processing environment (Lander et al. 2009) for real-time feedback during data collection. Images were collected on a Talos Arctica transmission electron microscope (Thermo Fisher) operating at 200 keV with a gun lens of 6, a spot size of 6, 70 μm C2 aperture and 100 μm objective aperture. Movies were collected using a K2 direct electron detector (Gatan Inc.) operating in counting mode at 45,000x corresponding to a physical pixel size of 0.91 Å/pixel. The dose rate was 4.413 e/pix/sec for a 10 second exposure, which makes for a total dose of 44.13 e/Å^2^ for the 1118 images collected at a defocus range of 0.8-2 μm.

##### 3D refinement

After the preprocessing pipeline, 425,087 particles were re-extracted to a pixel size of 1.22 Å/pixel with a box size of 271 Å. Using EMD-8743 as an initial model, we performed a 3D refinement in RELION-v3.0 (Zivanov et al. 2018) using D2 symmetry to obtain a structure at 3.2 Å resolution and B-factor of −110 Å^2^.

#### P-Rex1 screening

P-Rex1 samples were prepared as described (Cash et al. 2019) with the exception of details described in **Table 4**.

**Table 4.** Details of the automatic assessment of multiple P-Rex1 cryo-EM data sets.

|  | *18sep06b* | *18sep28b* | *18jan09b* | *18jan10d* | *18jul14a* | *18jan18c* |
| --- | --- | --- | --- | --- | --- | --- |
| **P-Rex1 concentration (μM)** | 3.0 | 3.0 | 3.0 | 3.0 | 3.0 | 3.0 |
| **Additive (μM)** | - | - | DDM (80) | Lubrol (40) | DDM (80) | DDM (80) |
| **Gβγ concentration (μM)** | - | - | - | - | 60 | 6.0 |
| **Grid type** | Quantifoil 1.2/1.3 | Lacey carbon | Quantifoil 1.2/1.3 | Quantifoil 1.2/1.3 | Quantifoil 1.2/1.3 | Quantifoil 1.2/1.3 |
| **Microscope** | Titan Krios | Talos Arctica | Talos Arctica | Talos Arctica | Titan Krios | Titan Krios |
| **Original Pixel Size (Å)** | 1 | 0.91 | 0.91 | 0.91 | 1 | 1 |
| **Number of total micrographs** | 1,716 | 1,491 | 1,206 | 1,110 | 1,352 | 5,011 |
| **Number of good micrographs #** | 986 | 445 | 841 | 790 | 1,217 | 4,157 |
| **Estimated diameter (Å)** | 144 | 144 | 135 | 132 | 138 | 151 |
| **Number of total picked particles** | 178,483 | 37,946 | 177,086 | 205,682 | 424,213 | 921,403 |
| **Number of good particles** | 94,514 | 9,535 | 114,630 | 141,982 | 145,941 | 492,883 |
| **Pixel size for 2D classification (Å)** | 4 | 3.59 | 3.64 | 3.67 | 3.94 | 3.97 |
| **Best diameter for 2D classification (Å)** | 129 | 216 | 108 | 105 | 110 | 120 |

**Table 5.** Overview of cryo-EM structures.

|  | **T20S Proteosome**<br>**(EMPIAR-10025)** | **HA Trimer**<br>**(EMPIAR-10175)** | **Aldolase** |
| --- | --- | --- | --- |
| **Microscope** | Titan Krios | Titan Krios | Talos Arctica |
| **Detector** | Gatan K2 | Gatan K2 | Gatan K2 |
| **Voltage (kV)** | 300 | 300 | 200 |
| **Electron exposure (e<sup>-</sup>/Å<sup>2</sup>)** | 53 | 73.24 | 44.13 |
| **Defocus range (µm)** | 0.9 - 2.4 | 1.0-2.1 | 0.8 - 2.0 |
| **Original pixel size (Å)** | 0.66 | 0.85 | 0.91 |
| **Symmetry imposed** | D7 | C3 | D2 |
| **Initial particle images**<br>**(no.)** | 52,153 | 167,788 | 536,520 |
| **Final pixel size** | 0.88 | 1.275 | 1.22 |
| **Final particle images (no.)** | 45,066 | 150,684 | 425,087 |
| **FSC threshold** | 0.143 | 0.143 | 0.143 |
| **Map resolution (Å)** | 3.1 | 3.2 | 3.2 |
| **B-factor (Å<sup>2</sup>)** | -103 | -151 | -110 |

---

**Figure S1.**
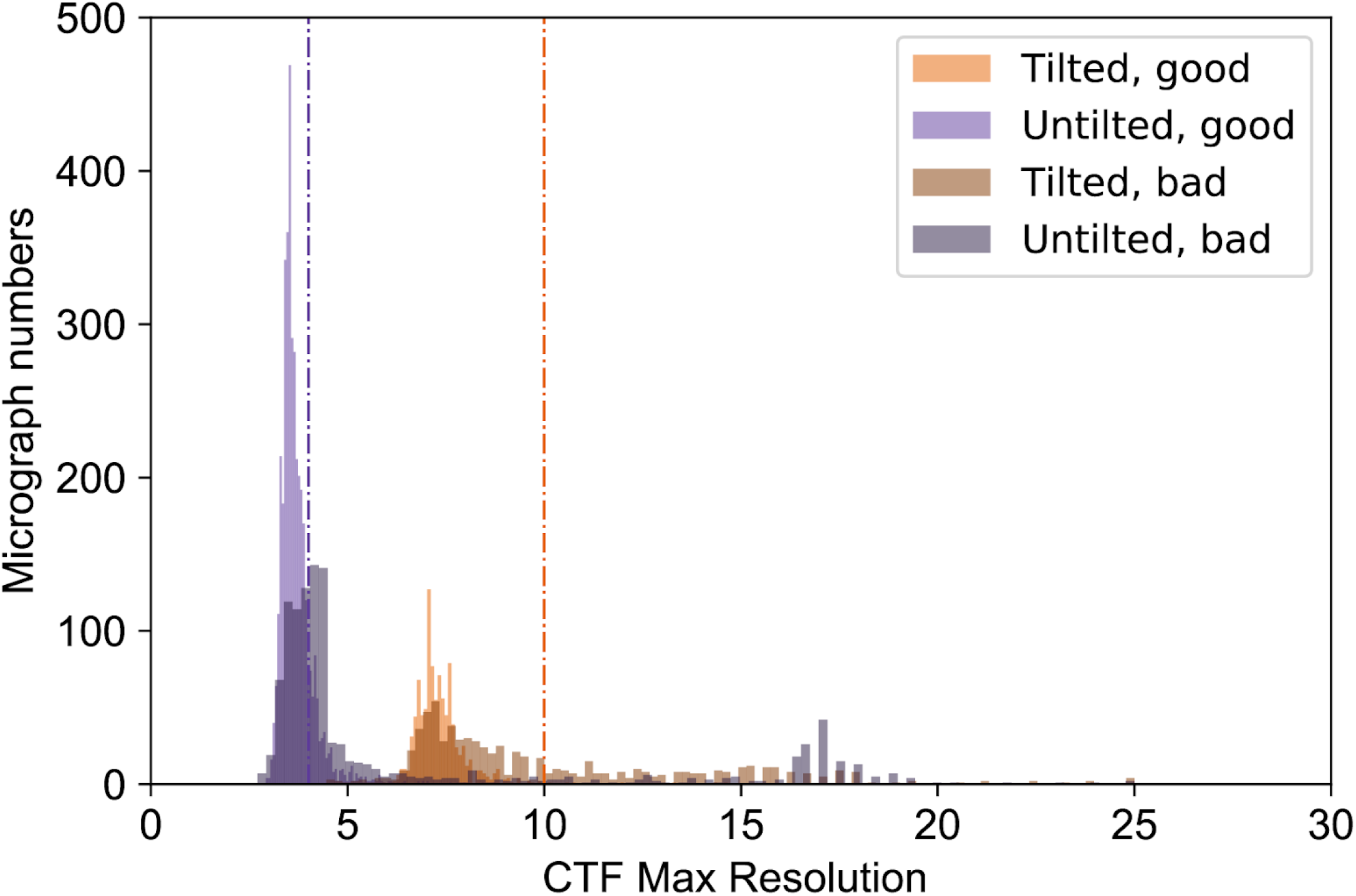
Comparison of CTF maximum resolution vs. ground truth (human assessment). Histograms of the CTF maximum resolutions outputted by CTFFIND4 of the P-Rex1 test set, color labeled according to the manually labeled ground truth. Vertical lines indicate 4 Å and 10 Å respectively.

**Figure S2.**
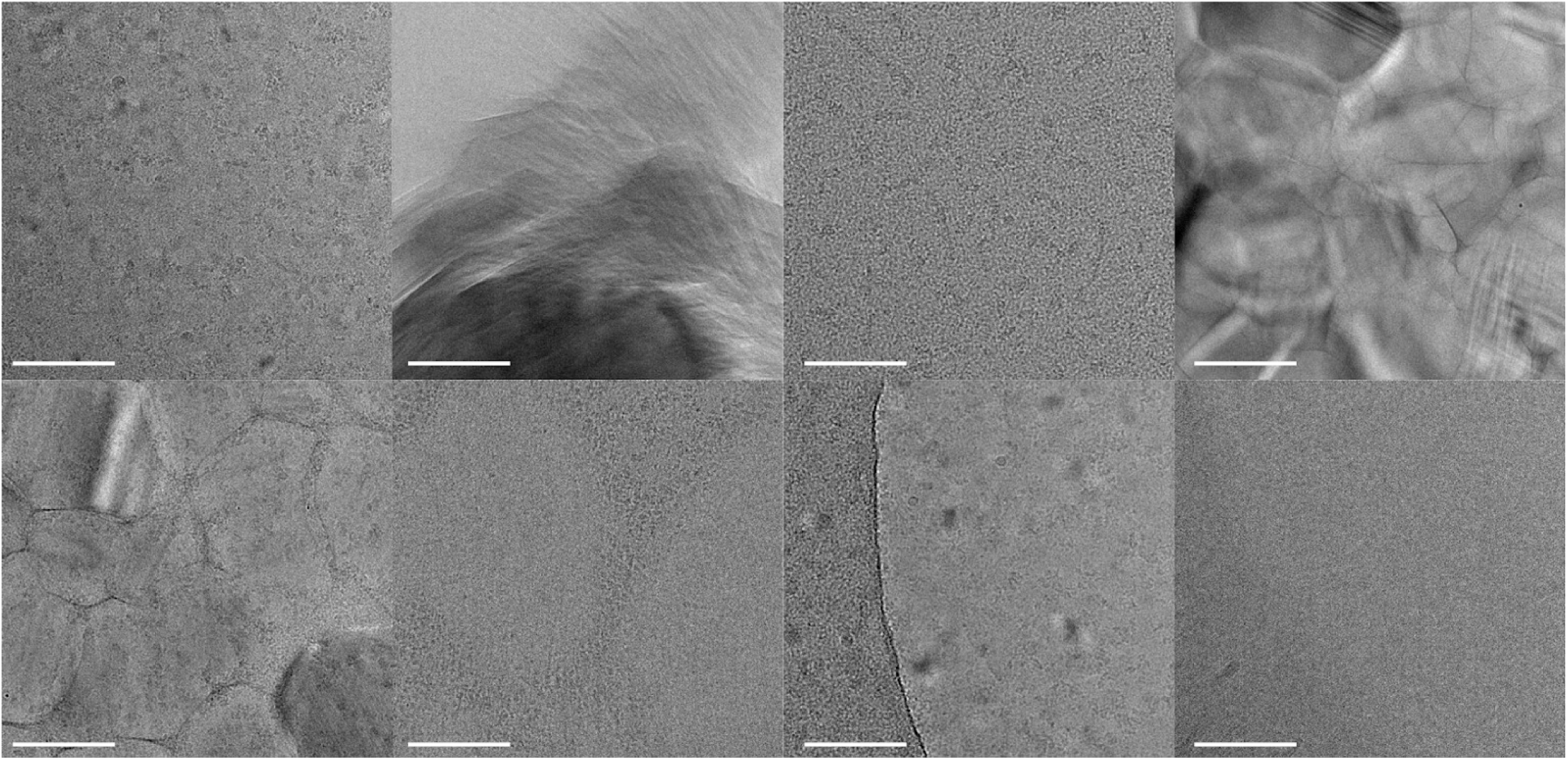
Examples of bad micrographs identified by *MicAssess* in the P-Rex1 test dataset.

**Figure S3.**
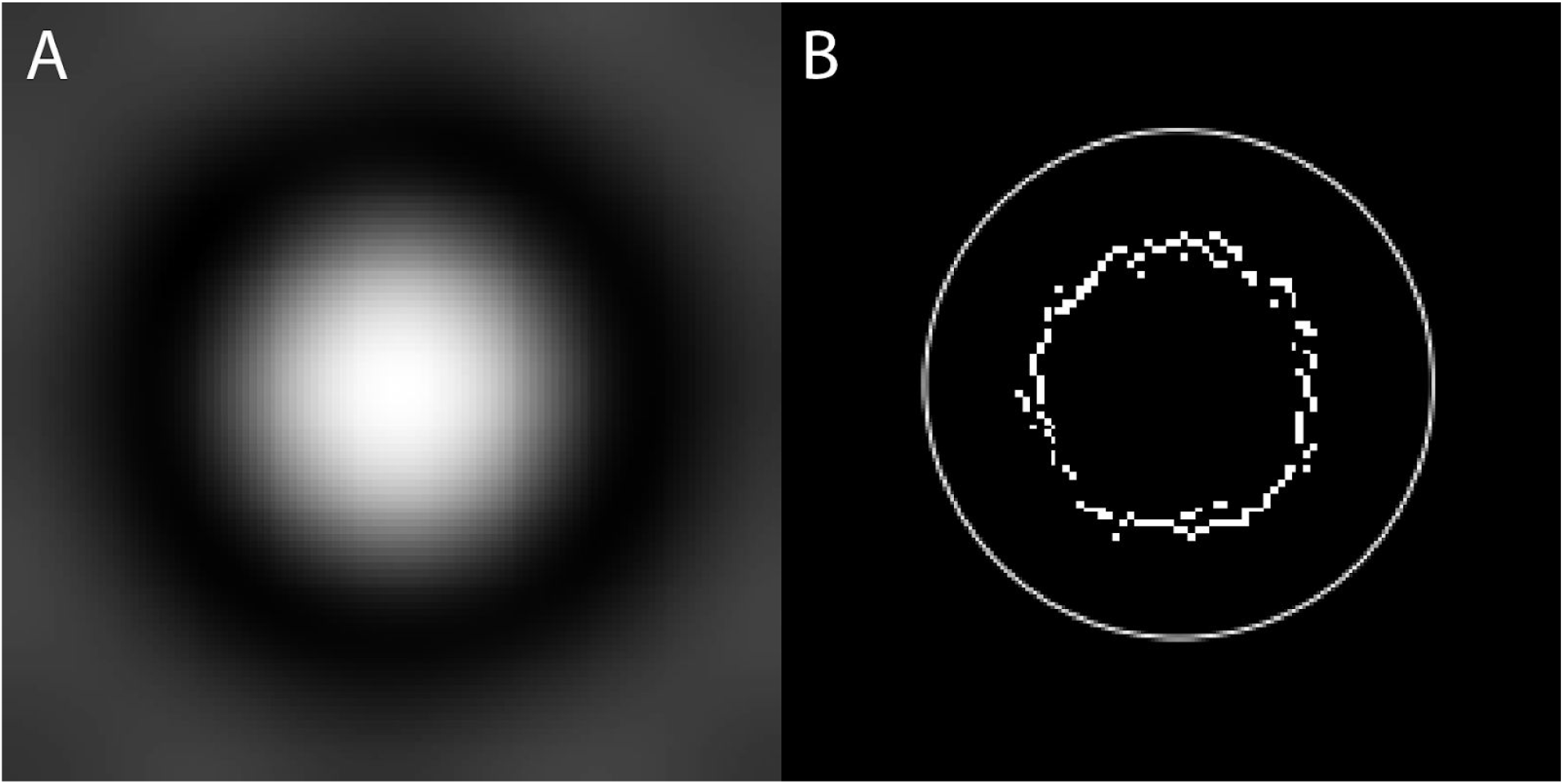
Particle size estimation. (A) Particles picked from 10 micrographs (EMPIAR-10025) were low pass filtered and averaged without any alignment. (B) Canny edge detector was applied to (A) and the edges found (inner circle) were dilated by a factor of 1.5 to estimate the diameter of the particle (outer circle).

**Figure S4.**
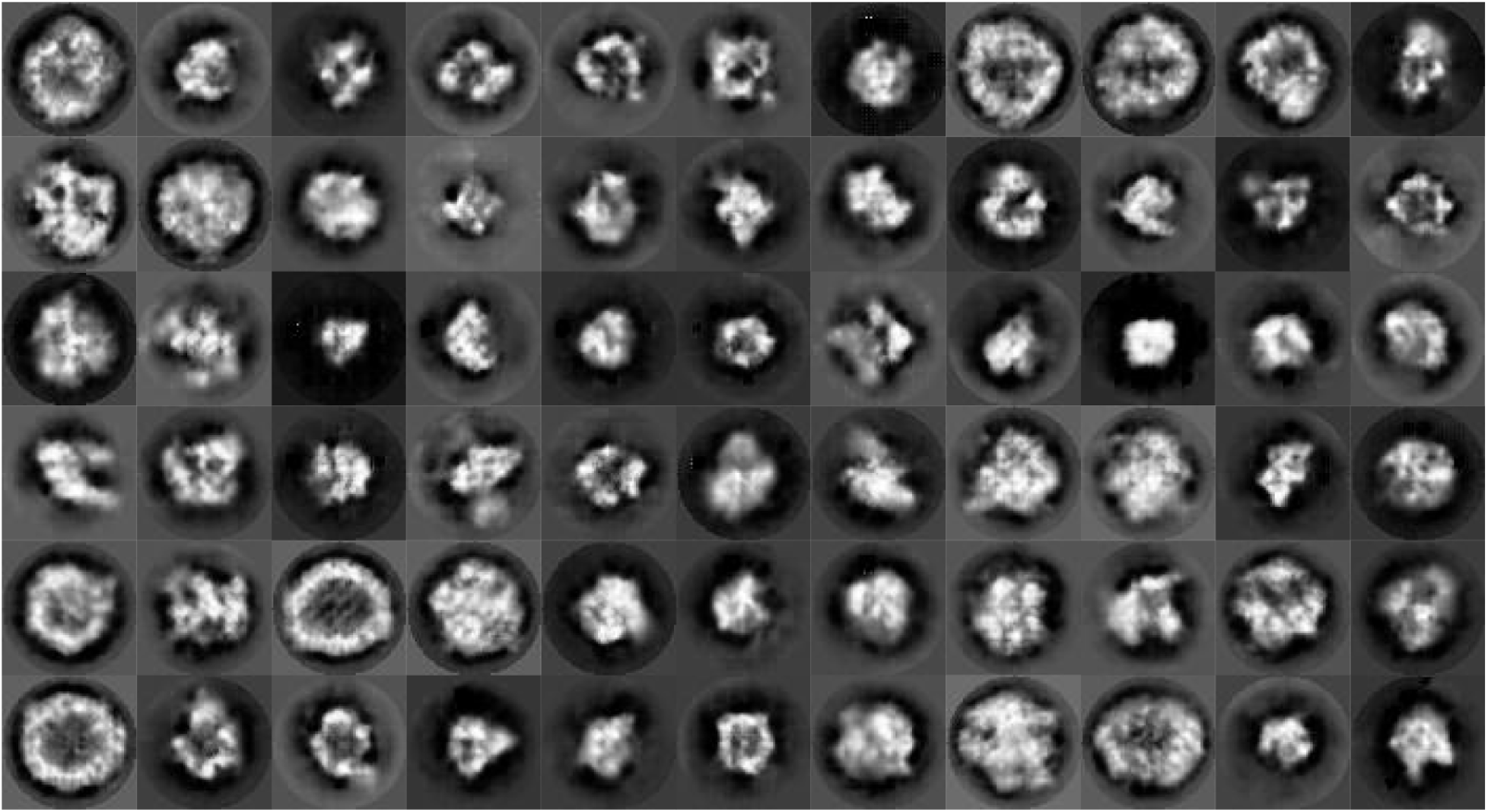
Artificial 2D class averages generated by DCGAN that were included in the training set of *2DAssess*.

**Figure S5.**
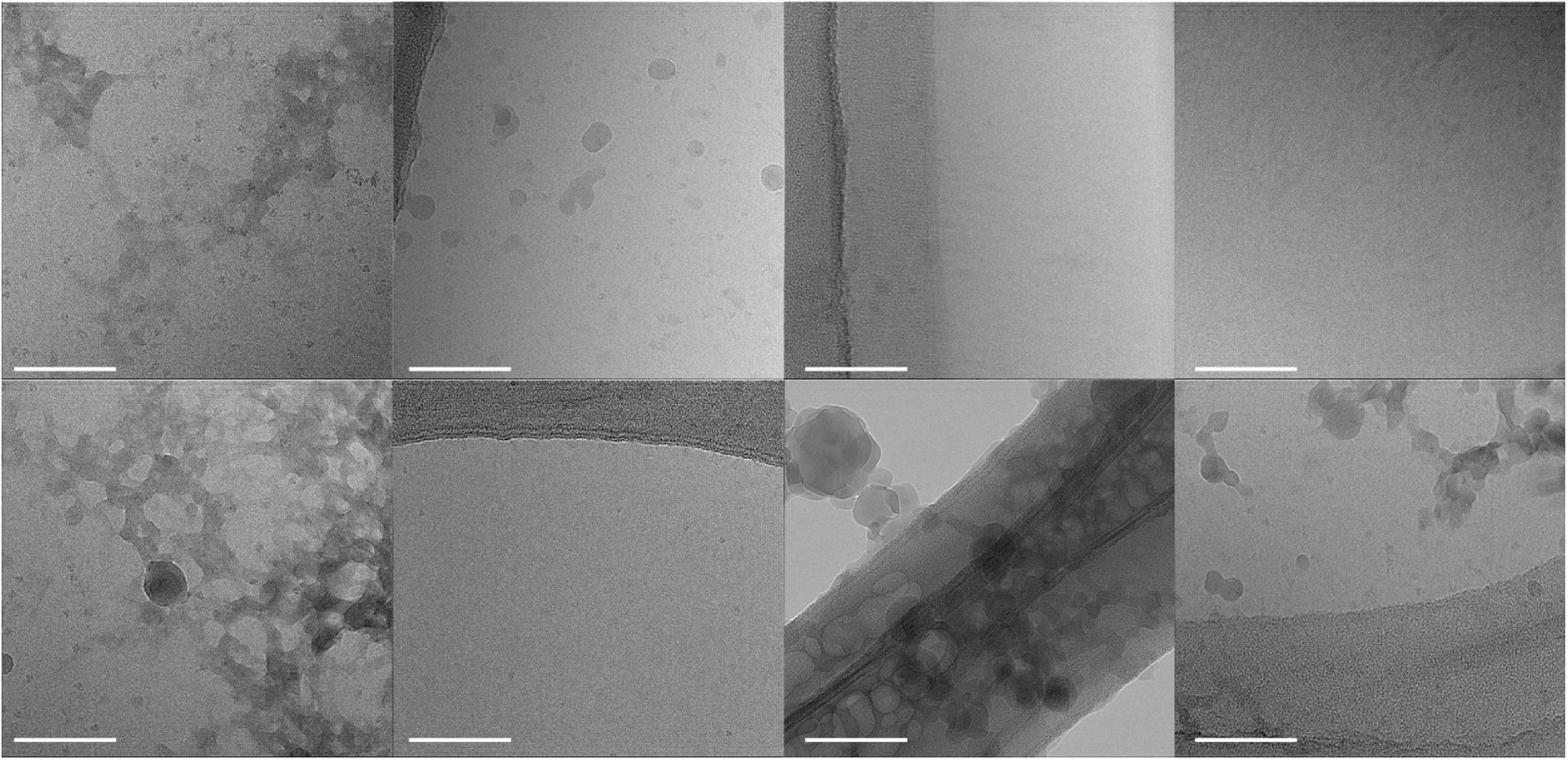
Examples of bad micrographs identified by *MicAssess* in the EMPIAR-10175 dataset. Scale bar = 100 nm.

**Figure S6.**
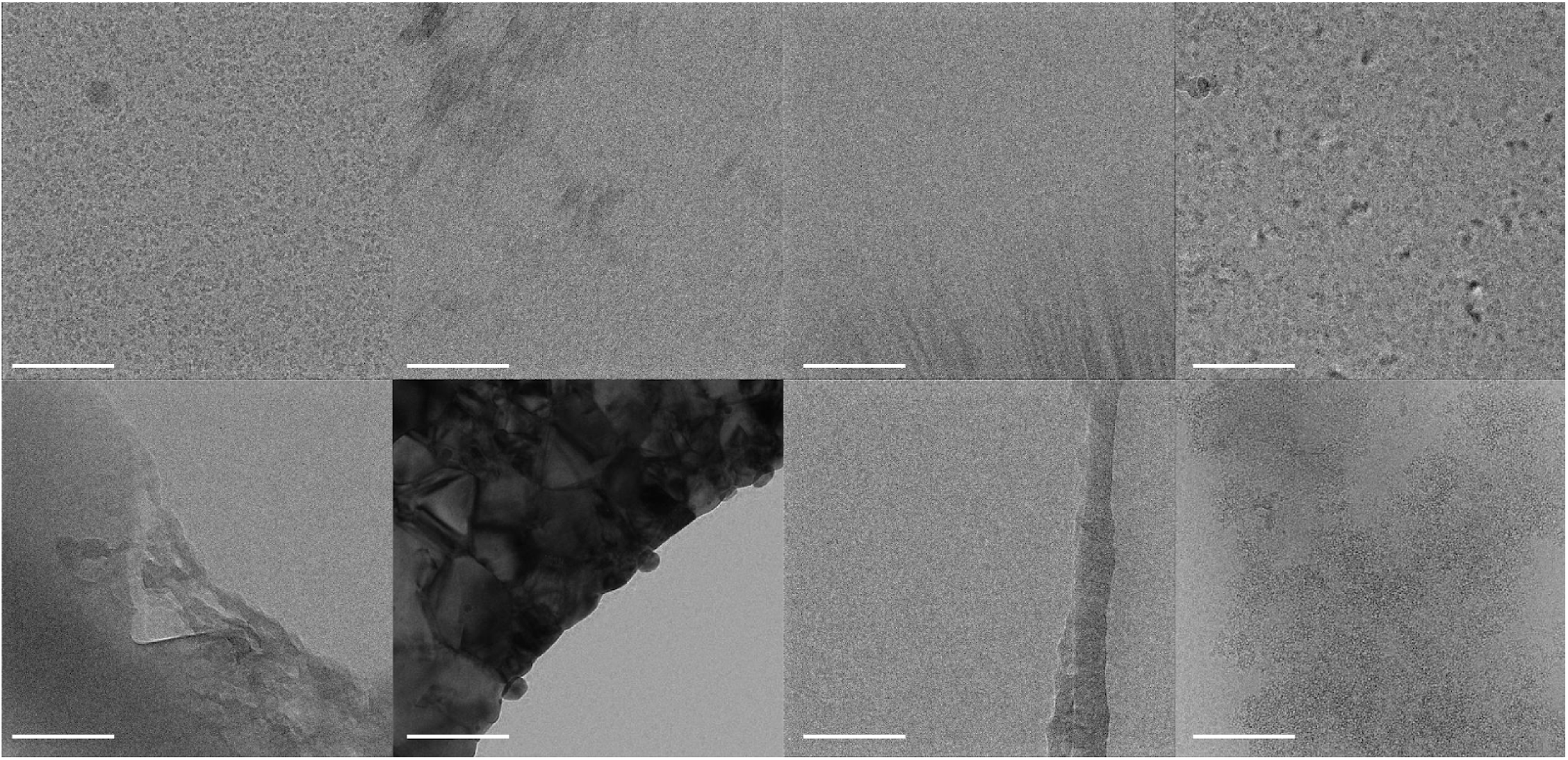
Examples of bad micrographs identified by *MicAssess* in the aldolase dataset. Scale bar = 100 nm.

**Figure S7.**
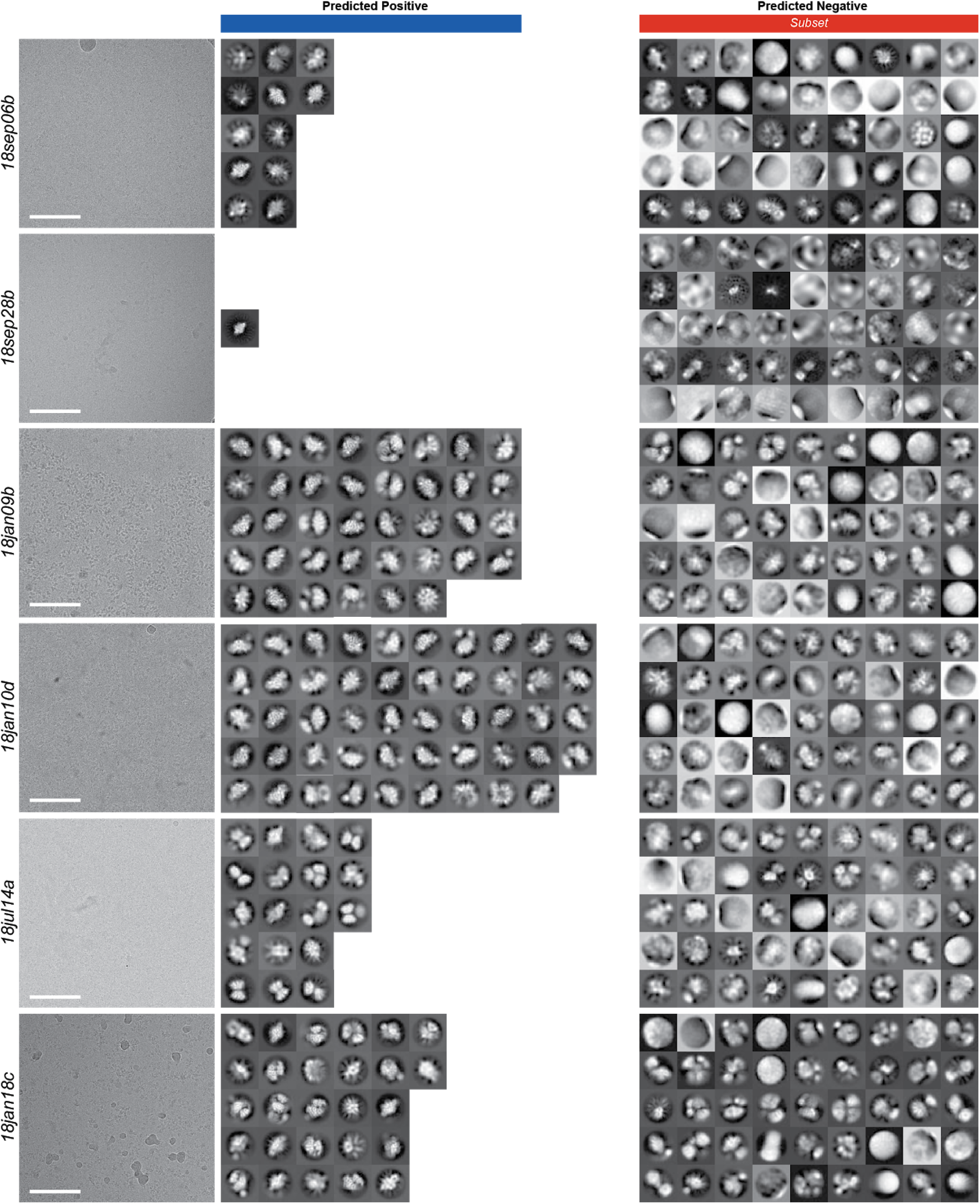
Example micrographs and 2D class averages from the automated analysis of P-Rex1 datasets. Showing all the good 2D class averages and a subset of the bad 2D class averages predicted by *2DAssess*. Scale bar in the micrographs = 100 nm.

